# Comparison of histological procedures and antigenicity of human post-mortem brains fixed with solutions used in gross anatomy laboratories

**DOI:** 10.1101/2024.01.18.576301

**Authors:** Eve-Marie Frigon, Amy Gérin-Lajoie, Mahsa Dadar, Denis Boire, Josefina Maranzano

## Abstract

**Background:** Brain banks provide small tissue samples to researchers, while gross anatomy laboratories could provide larger samples, including complete brains to neuroscientists. However, they are preserved with solutions appropriate for gross-dissection, different from the classic neutral-buffered formalin (NBF) used in brain banks. Our previous work in mice showed that two gross-anatomy laboratory solutions, a saturated-salt-solution (SSS) and an alcohol-formaldehyde-solution (AFS), preserve antigenicity of the main cellular markers. Our goal is now to compare the quality of histology and antigenicity preservation of human brains fixed with NBF by immersion (practice of brain banks) vs. those fixed with a SSS and an AFS by whole body perfusion, practice of gross-anatomy laboratories.

**Methods:** We used a convenience sample of 42 brains (31 males, 11 females; 25-90 years old) fixed with NBF (N=12), SSS (N=13), and AFS (N=17). One cm^3^ tissue blocks were cut, cryoprotected, frozen and sliced into 40 μm sections. The four cell populations were labeled using immunohistochemistry (neuronal-nuclei (NeuN), glial-fibrillary-acidic-protein (GFAP), ionized-calcium-binding-adaptor-molecule1 (Iba1) and myelin-proteolipid-protein (PLP)). We qualitatively assessed antigenicity and cell distribution, and compared the ease of manipulation of the sections, the microscopic tissue quality, and the quality of common histochemical stains (e.g., Cresyl violet, Luxol fast blue, etc.) across solutions.

**Results:** Sections of SSS-fixed brains were more difficult to manipulate and showed poorer tissue quality than those from brains fixed with the other solutions. The four antigens were preserved, and cell labeling was more often homogenous in AFS-fixed specimens. NeuN and GFAP were not always present in the NBF and SSS samples. Some antigens were heterogeneously distributed in some specimens, independently of the fixative, but an antigen retrieval protocol successfully recovered them. Finally, the histochemical stains were of sufficient quality regardless of the fixative, although neurons were more often paler in SSS-fixed specimens.

**Conclusion:** Antigenicity was preserved in human brains fixed with solutions used in human gross-anatomy (albeit the poorer quality of SSS-fixed specimens). For some specific variables, histology quality was superior in AFS-fixed brains. Furthermore, we show the feasibility of frequently used histochemical stains. These results are promising for neuroscientists interested in using brain specimens from anatomy laboratories.

## 1. Introduction

Normal and pathological aging of the human brain are topics of extensive research in the neuroscientific community. However, *in vivo* research is limited since standard imaging techniques, such as magnetic resonance imaging (MRI), do not provide information on molecular and cellular changes in the brain (at the cellular level of microstructure). Therefore, histology remains the Gold Standard method to unveil cellular changes that directly reflect underlying physio-pathological mechanisms (Cardoso et al., 2014; den Bakker, 2017; Durand-Martel et al., 2010). Ideally, brain tissue biopsies and surgical resections could be used for histological analysis (such as seen in (Kishore et al., 2023) for example) and correlated with *in vivo* MRI, but these samples are rarely available for research given the invasiveness and risks associated with brain biopsies. Hence, post-mortem (i.e., *ex vivo*) research is the preferred methodological approach, as it provides a larger amount of cerebral tissue available for histology and study of the molecular and cellular brain alterations.

Brain banks typically provide small blocks or single slices of post-mortem tissue to researchers. However, gross anatomy laboratories could potentially be an additional source of post-mortem brain tissue by providing more numerous and larger brain tissue samples, and even complete brains. Currently, these brains are not used in neuroscientific research since they are preserved with solutions optimized for the preservation of bodies dedicated to gross anatomy dissection, that differ in composition from the classic neutral-buffered formalin solution (NBF) used in brain banks (Beach et al., 2015; Carlos et al., 2019; Vonsattel et al., 2008). Formaldehyde, along with alcohol, are the two most widely used chemicals for their antiseptic and antibacterial properties, and their capacity to reduce autolysis and therefore prevent tissue decomposition (Brenner, 2014; Fox et al., 1985; Martins-Costa et al., 2022; Musiał et al., 2016; Shetty et al., 2020). They also increase cross-linking of the proteins, affecting the antigens’ conformation (Beach et al., 1987; Birkl et al., 2016; Dawe et al., 2009; Puchtler & Meloan, 1985; van Essen et al., 2010). Additionally, formaldehyde hardens and distorts the tissue (Mouritzen Dam, 1979; Weisbecker, 2011). Furthermore, formaldehyde has been proven carcinogenic (Fisher, 1905; Frølich et al., 1984; Musiał et al., 2016; Raja & Sultana, 2012; Thavarajah et al., 2012) and consequently, this has led to the search for alternative solutions in anatomy laboratories to reduce formaldehyde’s harmful effects.

For instance, anatomy departments, such as ours at the University of Quebec in Trois-Rivières, have developed and use other fixative solutions that combine various chemicals, such as a saturated salt solution (SSS) (Coleman & Kogan, 1998; Hayashi et al., 2016), and an alcohol-formaldehyde solution (AFS), which contain ethanol, phenol, glycerol, and isopropylic alcohol, and a minimal concentration of formaldehyde. Comparative studies across these new fixative solutions have been exclusively anatomical, i.e. focused on the tissue flexibility and color, ease of dissection, lifespan of the specimens after dissection (i.e. time before contamination or decomposition), health hazards, costs, etc. (Coleman & Kogan, 1998; Gulati et al., 2020; Hayashi et al., 2014; Martins-Costa et al., 2022; Shetty et al., 2020). However, to our knowledge, there are no studies on the ability of these solutions to preserve brain tissue adequately for (immuno)-histochemistry procedures. Instead, previous studies have focused on MRI (Benet et al., 2014; Kanawaku et al., 2014; Maranzano et al., 2020; Nazemorroaya et al., 2022; Schramek et al., 2013), other organs (Cabrera et al., 2017; Hammer et al., 2012; Hopwood et al., 1989; Lombardero et al., 2017; Rahman et al., 2021), or solutions with different chemical compositions than the SSS and the AFS currently employed by our laboratory (Hammer et al., 2012; Rahman et al., 2021; Tomalty et al., 2019). Therefore, these studies were limited with regards to information on tissue quality and antigen preservation in brain specimens fixed with such solutions, which is crucial to decide whether these brains are suitable for neuroscientific research.

Our previous work in mice showed that two solutions used in human gross anatomy laboratories (i.e., SSS and AFS), and the standard solution used in brain banks (i.e., NBF), preserved antigenicity of markers of the four main brain cell populations (i.e., neurons, astrocytes, microglia, and myelin of oligodendrocytes). Our work also showed that histological sections of brains fixed with SSS were difficult to manipulate and of poor quality but were still serviceable for immunohistochemical (IHC) and immunofluorescence (IF) analyses (Frigon et al., 2022). These promising results in mice might have been attributed to the lack of confounding variables that inevitably affect human samples (i.e., post-mortem delay, lengthy fixation period, variable ages of the donors, potential presence of vascular disease). Besides, human and mice brain characteristics are not alike (Ventura-Antunes et al., 2013), and as such, fixation quality through vascular perfusion of the specimens might not be comparable. Furthermore, as cellular densities differ (Herculano-Houzel et al., 2011; Loomba et al., 2022; Ventura-Antunes et al., 2013), we cannot deduce whether antigenicity and histological quality are preserved in human brain tissue in a similar fashion. Also, conclusive evidence on whether common histochemical stains, typically used for diagnostic purposes and neuropathology research, are feasible on brains fixed with solutions such as SSS or AFS is lacking. Therefore, several histological procedures need to be explored in post-mortem human brain tissue fixed with solutions typically used by anatomists.

The goal of the present study is to compare the quality of histological outcomes applied to human brains fixed using brain banks’ techniques (i.e., fixation by immersion in NBF) to those fixed with SSS or AFS by perfusion as applied in gross anatomy laboratories. We compared the ease of manipulation of the tissue, the tissue quality, the IHC labeling quality of four antigens of interest (neurons (anti-NeuN), astrocytes (anti-GFAP), microglia (anti-Iba1) and oligodendrocytic myelin protein (anti-PLP)) and the quality of various histochemical stains (i.e., Cresyl Violet, Luxol Fast Blue, Prussian blue and Bielchowsky’s) across the three experimental groups.

## 2. Material and methods

### 2.1 Population

We used a convenience sample of 42 human brains (NBF: N=12; SSS: N=13; AFS: N=17) from our body donation program (Université du Québec à Trois-Rivières, Canada) as well as the Douglas-Bell Canada Brain Bank (Douglas Mental Health University Institute, McGill University, Montréal, Canada). Prior to their death, the donors to the anatomy laboratory provided consent on body donation and sharing of medical information for teaching and/or research purposes. The study was approved by the University’s Ethics Subcommittee on Anatomical Teaching and Research. Donors from the Brain Bank were obtained in collaboration with the Quebec Coroner’s Office and with informed consent from the nearest relative, following the guidelines of the Douglas Research Centre Ethic Committee.

The median age at time of death of the donors was 76 years old (range: 25 to 90). The male to female ratio was 3:1. The case number, age, sex, cause of death, post-mortem delay to the fixation treatment (i.e., post-mortem interval) and delay until the histology processing of the tissue, are reported in table 1.

**Table 1.**
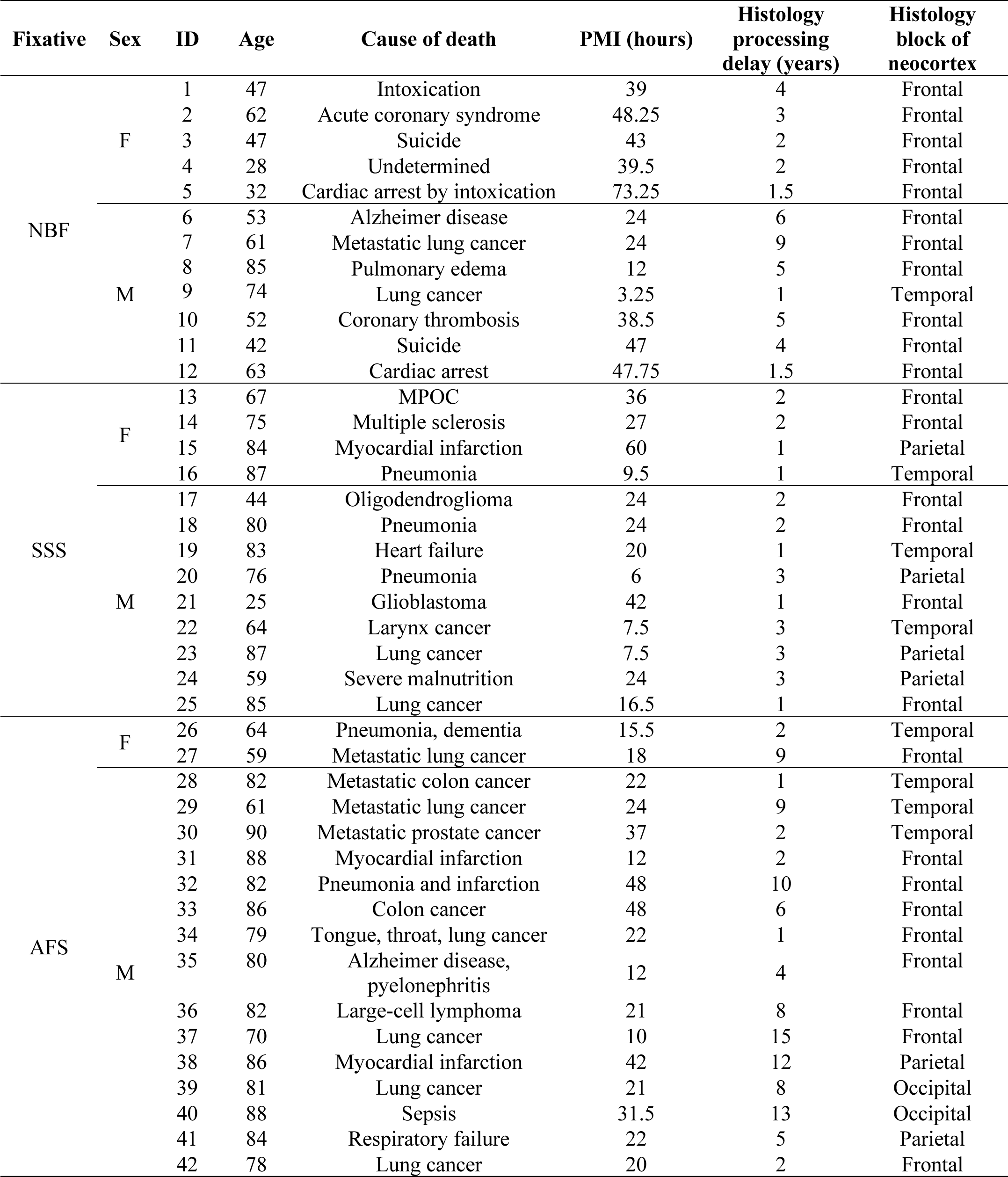
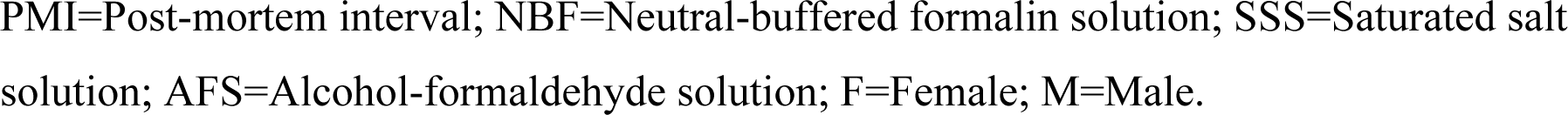
Specimen data.

### 2.2 Fixation procedure

All specimens were fixed with one of three solutions (chemical components are shown in Table 2). NBF-fixed brains were extracted fresh and then immersed in 4 liters of NBF following the brain bank technique and kept in formaldehyde until use. For SSS and AFS, the bodies were perfused with 25 L of one of the solutions through the common carotid artery using a pump (41 kPa) (Duotronic III, Hygeco International Products) which is the common embalming technique performed in gross anatomy laboratories (Maranzano et al., 2020). The brains were extracted, and 1 cm^3^ blocks (neocortex for gray matter (GM) and subcortical white matter (WM)) were taken, choosing an area that was macroscopically of good quality (no superficial tears, good color and texture) and not damaged during the extraction and arachnoid removal procedure.

**Table 2.**
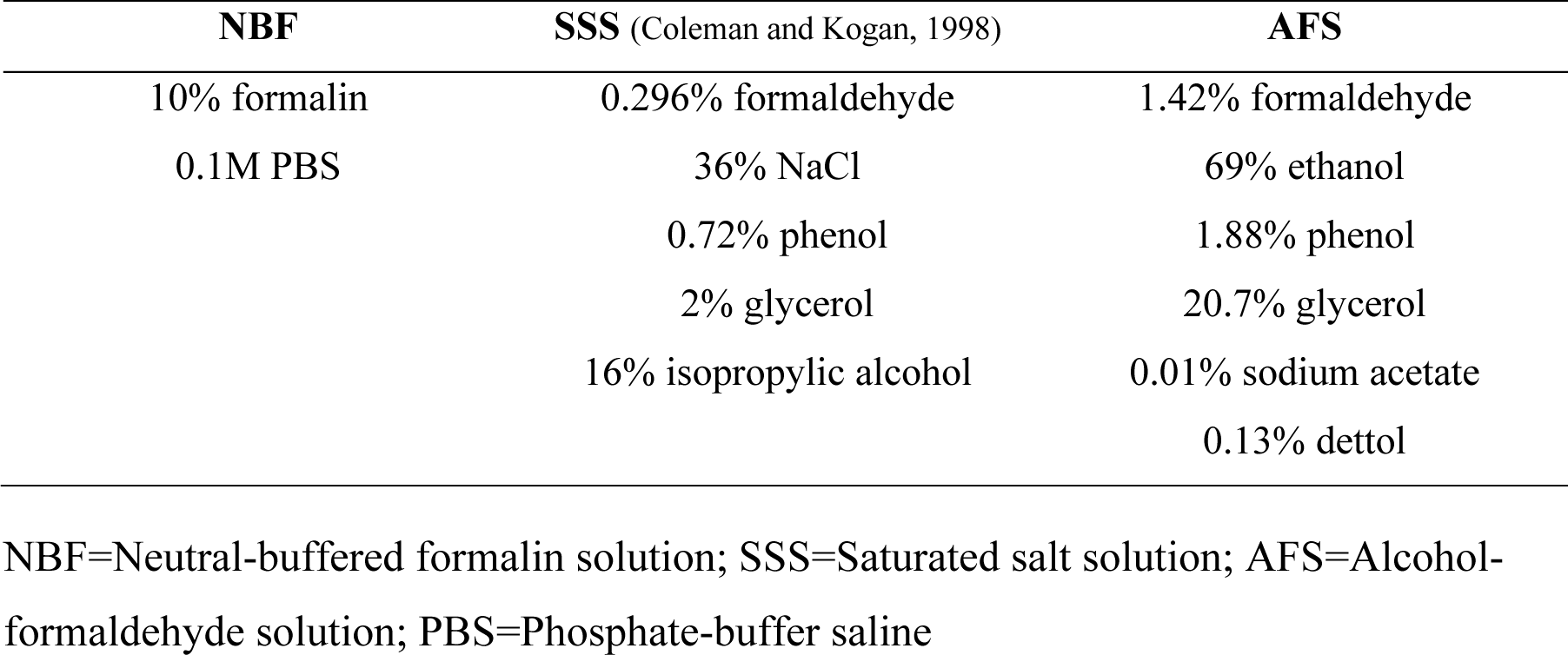
Fixative’s components.

| NBF | SSS (Coleman and Kogan, 1998) | AFS |
| --- | --- | --- |
| 10% formalin | 0.296% formaldehyde | 1.42% formaldehyde |
| 0.1M PBS | 36% NaCl | 69% ethanol |
|  | 0.72% phenol | 1.88% phenol |
|  | 2% glycerol | 20.7% glycerol |
|  | 16% isopropyl alcohol | 0.01% sodium acetate |
|  |  | 0.13% dettol |

NBF=Neutral-buffered formalin solution; SSS=Saturated salt solution; AFS=Alcohol-formaldehyde solution; PBS=Phosphate-buffer saline

### 2.3 Histology processing

Blocks were rinsed in 0.1M PBS and immersed in 30% sucrose for 2 to 5 days (until the floating block sinks, which is the sign of appropriate sucrose diffusion into the tissue) for cryoprotection. Blocks were then frozen on dry ice before cutting 40 μm sections on a cryostat (Leica CM1950) at −19°C, that were either 1-mounted on 2% gelatin-subbed slides for the histochemical stains (Cresyl violet, Luxol Fast blue, Prussian blue and Bielchowsky’s), or 2-placed in well plate with PBS for subsequent free-floating IHC procedures. After IHC or histochemical staining (see the sections below), the sections were dehydrated in 5-minute baths of increasing grades of ethanol (70%, 95%, 100%) and coverslipped using Eukitt.

### 2.4 Immunohistochemistry

Free-floating sections were rinsed three times for five minutes in 0.1M PBS. They were then incubated in a 20% aqueous methanol solution with 0.5% H_2_O_2_ for thirty minutes to quench endogenous peroxidase. After another round of rinses (3×5minutes in 0.1M PBS), sections were incubated in blocking solution for two hours (3% Normal Donkey Serum; 0.5% Bovine Serum Albumin; 0.3% Triton X-100 in 0.1M PBS), followed by the primary antibodies’ (Table 3) incubation at 4°C in the same blocking solution overnight. Sections were rinsed again (3×5minutes in 0.1M PBS) before a two-hour incubation in donkey anti-rabbit biotinylated secondary antibody (1:500) at room temperature in the same blocking solution. After another rinse, sections were incubated in an avidin-biotin complex (ABC) kit for thirty minutes in the dark (VECTASTAIN® Elite® ABC-HRP Kit, Vector Laboratories, catalog#VECTPK6100). Sections were then rinsed for five minutes in TRIS-buffered saline (TBS) (0.05M; 0.9% NaCl) before the final incubation in 0.07% diaminobenzidine (DAB) (Sigma, #D5905-50TAB) with 0.024% H_2_O_2_ in TBS for ten minutes.

**Table 3.**
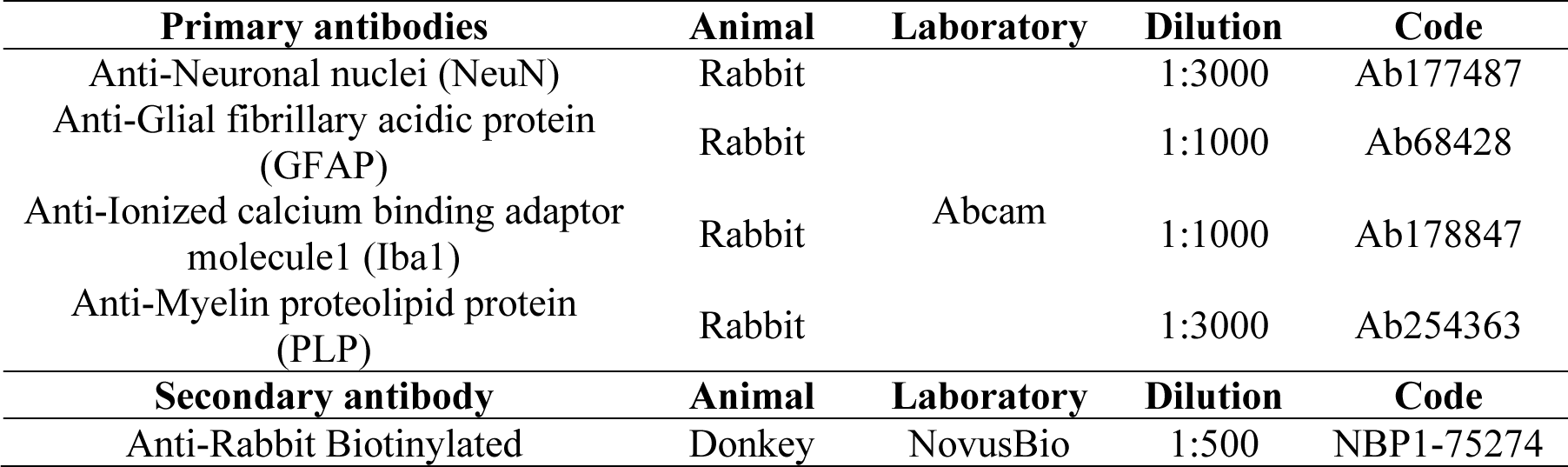
Immunohistochemical antibodies.

### 2.5 Histochemical stains

All specimens were also processed for four histological stains, commonly used in brain histopathology, namely 1-Cresyl Violet (endoplasmic reticulum of the cell nuclei, Nissl bodies), 2-Luxol Fast Blue (myelin), 3-Prussian blue (iron stain with a neutral red counterstain for Nissl bodies), and 4-Bielchowsky’s (neurofibrils and axons). Prior to the staining, sections were defatted in Xylene for 30 minutes and hydrated in increasing alcohols −100% Ethanol 2 × 1 minute; 3-95% Ethanol 2 × 1 minute; 70% Ethanol 2 × 1 minute; H_2_O_d_ 1 minute. After the staining (see protocols below), a dehydration process was applied using the same baths, but in a reverse order.

#### 2.5.1 Cresyl Violet

Cresyl violet (CV) stains the endoplasmic reticulum and the cellular nuclei in dark purple, which includes, in the brain, cell bodies of neurons and glial cells (Eilam-Altstädter et al., 2021). The protocol followed guidelines of US Army Inst Pathol (Luna et al., 1968), in which the sections were hydrated first, submerged for 1h15 in 0.01% Cresyl violet (Sigma Aldrich catalog #C5042) in a 0.1M acetate buffer (Anachemia, AC-8218) and 0.1M acetic acid (BDH 10001CU), and then dehydrated.

#### 2.5.2 Luxol Fast Blue

Luxol flast blue (LFB) stains myelin, and should be dark blue in the WM, and lighter blue (or even clear) in the GM. We followed principal guidelines and adapted the protocol from Pathology Center-Histological methods for CNS (https://pathologycenter.jp/method-e/) (Tokyo Metropolitan Institute of Medical Science) for this staining. The sections were incubated 2 hours at 45°C in 0.1% Luxol Fast Blue (Solvent blue 38, Sigma Aldrich, catalog #S3352) in 95% Ethanol, then rinsed in H_2_O_d_. The differentiation (clearing over-dye) was performed in 0.05% Lithium carbonate (Sigma Aldrich, catalog #62470) for 1 minute before immersion in 70% Ethanol for 10 minutes.

#### 2.5.3 Prussian Blue

Prussian blue was performed to stain ferric iron in blue in combination with a Nuclear fast red counterstain which stains cell nuclei in pink. Iron Stain Kit (Abcam, ab150674) was used according to the manufacturer’s protocol (https://www.abcam.com/products/assay-kits/iron-stain-kit-prussian-blue-stain-ab150674.html#). In order to assess the feasibility of this staining, we included cases diagnosed with Alzheimer’s disease in which we expected to detect iron (LeVine, 1997). Once validated, we proceeded to stain all the other specimens.

#### 2.5.4 Bielchowsky silver stain

Bielchowsky’s method is a silver staining protocol to label nerve fibers and neurofibrils in dark brown/black, and is commonly used to demonstrate neurofibrillary tangles and senile plaques in the central nervous system (Stahnisch, 2015). Our protocol was inspired from Bielchowsky’s Silver Stain protocol by Paul Polak and Douglas Feinstein, University of Illinois, Chicago (abcam) and IHC world (https://www.ihcworld.com/_protocols/special_stains/bielschowsky.htm) and adapted as follows: sections were placed into a 10% Silver nitrate (Sigma Aldrich, catalog #209139) solution for 20 minutes at 40°C, following 3 rinses in water and another 20 minutes incubation at 40°C in the same silver nitrate solution that was titrated with high-concentrated ammonium hydroxide (ThermoScientific, catalog #423305000). Sections were then placed 1 minute in a working developer solution (8 drops of Stock Solution + 8 drops of ammonium hydroxide + 50 ml of water; Stock solution: 20 ml of 37% formaldehyde + 0.5g of Citric acid + 2 drops of Nitric acid + 100 ml of water), followed by a 1% ammonium hydroxide incubation. Sections were rinsed in water three times before a 5 min bath of 5% Sodium Thiosulfate (Sigma Aldrich, catalog #S7026).

### 2.6 Variables of interest

The sections were assessed for multiple variables using a brightfield microscope (Olympus BX51W1) controlled by Neurolucida software (MBF Bioscience). We scored the IHC and histochemical stains variables to create qualitative categories that were compared for the three groups (fixatives) by looking through all the sections that were available for each specific staining. All variables were scored by a rater having 5-years experience in histology procedures and microscopy analysis (EMF). Figure 1 shows scoring of the criteria for different IHC variables (tissue quality and antigenicity preservation) that were assessed, while Figure 2 shows scoring of the criteria used for the four histochemical stains.

**Figure 1:**
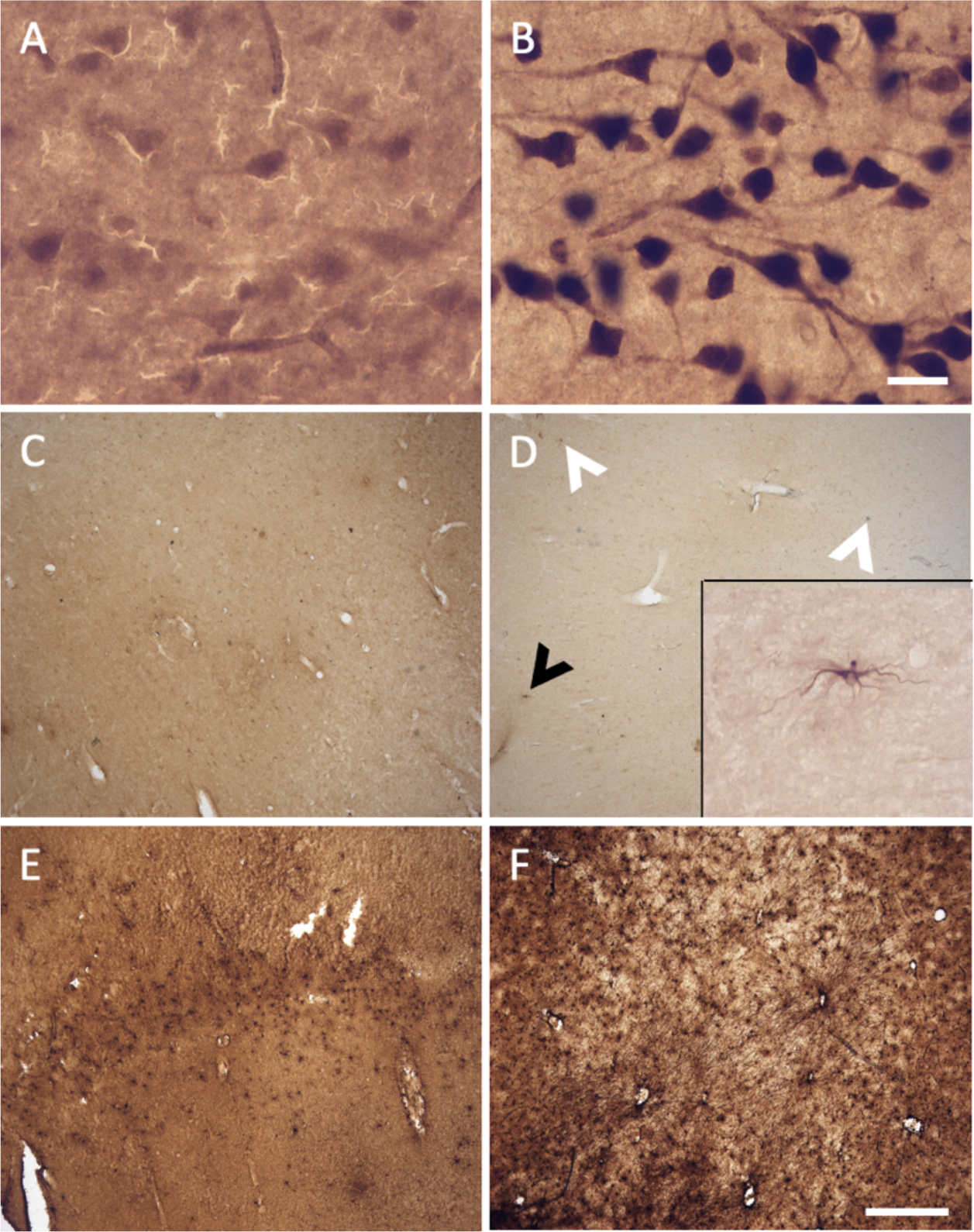
Photomicrographs of different criteria of the IHC categorical variables that were assessed. Photomicrographs (60X) of a fissured neuropil and shrunken and shriveled cells (A), compared to a uniform neuropil with regular cell contour (B) (Section 2.6.2.1 and 2.6.2.2). Scale bar = 25 μm (valid for A and B). Photomicrographs (4X) of the antigenicity preservation and distribution as an example using GFAP labeling, where we can see astrocytes that were completely absent (C); isolated (D) (where arrowheads point out the cells, the black one being shown in the 60X insert; in patches (E) or homogeneously distributed through the full analyzed sections (F) (Section 2.6.3). Scale bar = 500 μm (valid for C-F).

**Figure 2:**
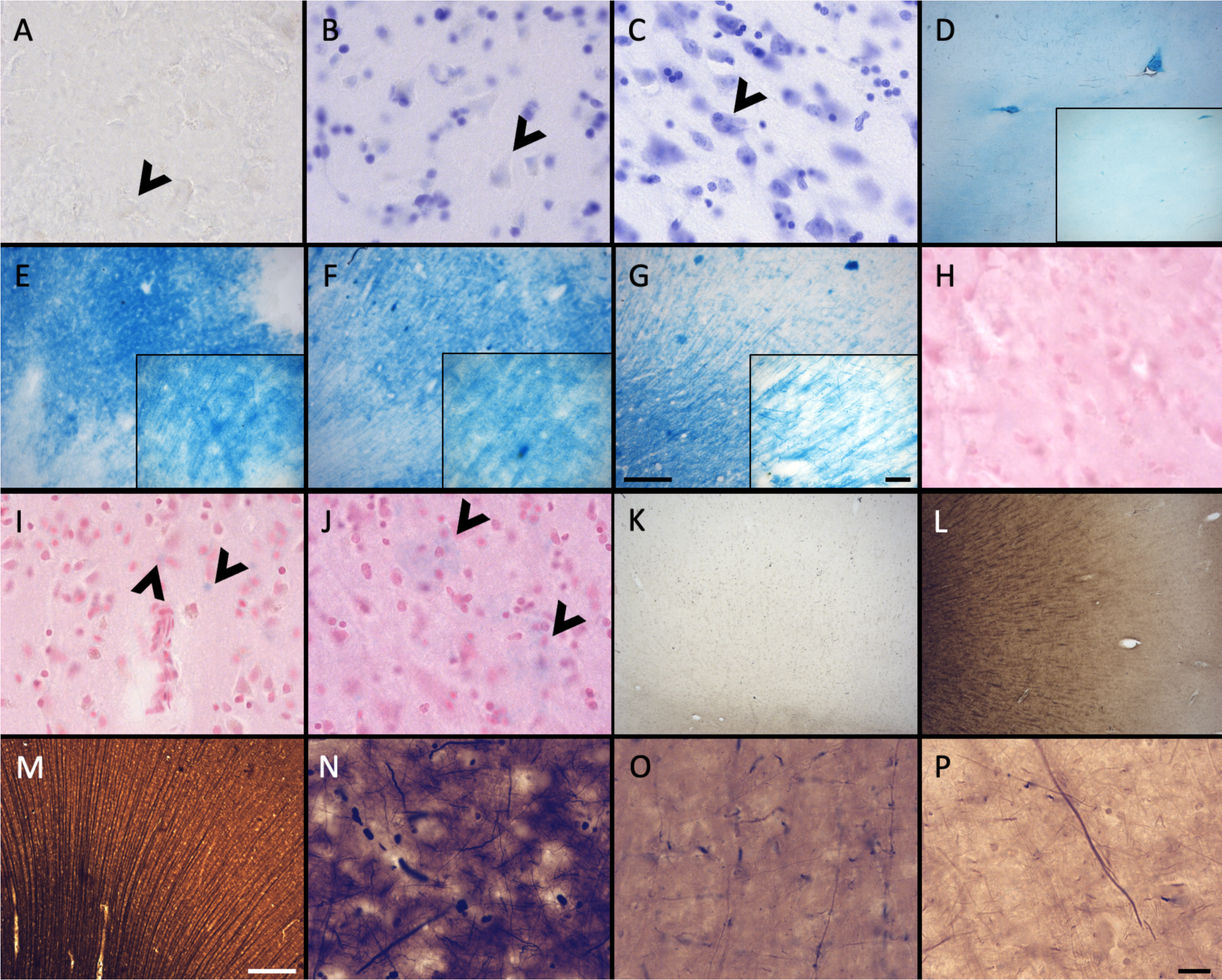
Photomicrographs of different criteria of the histochemical stains’ categorical variables that were assessed. Photomicrographs (60X) of an absent cresyl violet staining (A), compared to when the neurons appear pale (B), or strongly labeled (dark) neurons (C) (Section 2.6.5.1). Arrowheads point out neurons. Photomicrographs (4X) of the distribution of fibers and WM-GM contrasts of D) no fibers and an heterogenous blue stains; E) heterogenous fibers and differentiation; F) homogenous fibers, but bad differentiation; or G) homogenous fibers and good differentiation. Scale bar= 500 μm (valid for D-G). Inserts in D to G show photomicrographs (20X) of the myelin fibers in the respective categories (Section 2.6.5.2). Scale bar = 100 μm (valid for D-G inserts). Photomicrographs (60X) of an absent neutral red (counterstain of the Prussian blue) staining (H), compared to when the neurons appear pale (I), or strongly labeled (dark) neurons (J) (Section 2.6.5.3). Arrowheads show blue spots, representing iron accumulation. Photomicrographs (4X) of the fibers preservation in the Bielchowsky’s stain, where we can see absence of fibers (K), heterogenous labeling of fiber (L) or homogenous distribution of fibers (M). Scale bar = 500 μm (valid for K-M). Photomicrographs (60X) of the background in Bielchowsky’s stain that could be affected by silver deposits and therefore be dark (N), intermediate (O), or light (P). Scale bar = 25 μm (valid for A-C, H-J and N-P).

#### 2.6.1 Ease of manipulation

The first variable of interest was assessed during the processing of sections, i.e., cutting, well transfers, mounting on slides. We used four signs to assess the ease of manipulation: 1-slice consistency (soft or firm), 2-presence of tears (yes/no), 3-slices sticking to the brush that was used to manipulate/mount the slice (yes/no), and 4-presence of rolling slices (yes/no). Scores ranged from: Very poor (4 signs present=0), Poor (3 signs present=1), Good (1 or 2 signs present=2) to Very good (No sign present=3).

#### 2.6.2 Tissue quality

We used two criteria to assess this variable, namely the neuropil uniformity and the cellular shape, since it reflects the integrity of the tissue and the cellular morphology, susceptible to be differentially affected by the different chemicals (Spencer, 2017; Troiano et al., 2009) and the histology delay, which was in some specimens longer than five years. These criteria were assessed in the NeuN stained sections by scanning through the thickness of sections with a 60X objective (60X, UPlanSApo 60x/1.40 Oil∞/0.17/FN26.5 UIS2).

##### Neuropil uniformity

The tissue was of lower quality when there was presence of a fissured neuropil. Scores indicated: Fissured=0 (Figure 1A), Uniform=1 (Figure 1B).

##### Cellular shape

The neurons showed signs of altered shape (i.e., lower quality) if they presented an irregular and shriveled contour (irregular). Scores indicated: Irregular (shrunken/shriveled)=0 (Figure 1A), Regular (smooth)=1 (Figure 1B).

#### 2.6.3 Antigenicity preservation

We looked at sections targeted for all the antigens to verify the cell distribution across the section, using a 4X objective to have a large field of view (4X, UPlanSApo 4x/0.16∞/-/FN26.5). Antigenicity pres4ervation was scored from worse (reflecting destruction/cross-linking of the antigen) to best (reflecting ideal preservation of the antigen) as follows: 1-absence of labeling (Figure 1C); 2-heterogeneous distribution of the cells (presence of isolated cells (Figure 1D, E), 3-cells’ patches (Figure 1F) or 4-a homogeneous distribution (Figure 1G). This was assessed for all the immunolabeled sections of the four antigens of interest. Scores indicated: Absence=0 (Figure 1C), Isolated cells=1 (Figure 1D, E), Cell patches=2 (Figure 1F), Homogeneous distribution=3 (Figure 1G).

Specimens that showed no antigenicity for one or more antigens or heterogeneous distribution (i.e., isolated cells or cell patches) were reprocessed following an antigen retrieval protocol. To this end, sections were subjected to 20-minute boiling-citrate buffer (free-floating) and then immersed into a cold-water bath for 10 minutes. The IHC procedures were then performed following the same protocols (section 2.4), and the sections were analyzed anew.

#### 2.6.4 Histochemical stains’ quality

We assessed four histochemical stains to evaluate the preservation of different cell populations in a non-immunological way.

##### Cresyl Violet

A quality Nissl stain shows a clear contrast between well stained neuronal cell bodies with clear cytoplasmic granulations and the clear surrounding neuropil. Glial cells were strongly labeled in all cases, but neurons were heterogeneous in their staining. Therefore, a high contrast between the stained neuronal cell bodies and neuropil was considered as indicative of quality. This was assessed at low (4X) and high (60X) magnification of all the labeled sections available. Scores indicated: No staining=0 (Figure 2A), Pale neurons=1 (Figure 2B), Dark neurons=2 (Figure 2C).

##### Luxol Fast blue

Luxol fast blue staining is known to dye myelinated axons and the oligodendrocytes’ nuclei that produce this myelin sheath in the white matter of the brain (Lindberg & Lamps, 2018). LFB is frequently used in the diagnosis of multiple sclerosis, Alzheimer’s disease, or white matter hyperintensities (Bronge et al., 2002; Gouw et al., 2011; Humphreys et al., 2019; Kuhlmann et al., 2017; Laule et al., 2008; Laule & Moore, 2018; Ouellette et al., 2020; Roseborough et al., 2020; Selvaraji et al., 2022). This staining was therefore important to assess in brains fixed with other solutions. The LFB stain was assessed by the contrast between the WM and GM at low (4X) and high (60X) magnification. The WM should show homogeneous distribution of fibers and be dark blue and should be clearly differentiated from the GM that should be free of staining or light blue. Scores indicated: No fibers and heterogenous blue stain=0 (Figure 2D); Heterogenous fibers and differentiation=1 (Figure 2E), Homogenous fibers but bad differentiation=2 (Figure 2F) and Homogenous fibers and good differentiation=3 (Figure 2G).

##### Prussian blue

Prussian blue staining (Perl’s stain) is known to reveal ferric iron deposits in the neuropil of the brain (Churukian, 2008; Guindi, 2018; Xiao et al., 2023). This staining is of interest in neurodegenerative diseases, small vessel diseases, or white matter pathology detected by MRI, in which brains may be affected by such iron deposits (Connor & Benkovic, 1992; Dusek et al., 2022; Kruer, 2013; Roseborough et al., 2020). Since iron deposits did not occur in all specimens, this was not assessed in this staining; only the feasibility of the iron revelation in brains fixed with the three solutions was considered. However, we assessed the neutral red staining of the Prussian blue counterstain in a similar fashion as the cresyl violet. Scores indicated: No staining=0 (Figure 2H), Pale neurons=1 (Figure 2I), Dark neurons=2 (Figure 2J).

##### Bielchowsky’s

Bielchowsky’s silver stain is known as a silver impregnation method for nervous tissu (Ogawa, 1990) which is used to visualize axons, neurofibrils and neurofibrillary tangles or neuritic plaques (Allsop, 2000; Elobeid et al., 2014; Fan et al., 2018; Kovacs & Budka, 2010; Love & Nicoll, 1992; Switzer, 2000; Uchihara, 2007; Wisniewski et al., 1989). The quality was assessed using two criteria: the fibers preservation/distribution and the background intensity.

##### Fibers’ preservation and distribution

The stained sections were scanned at low (4X) magnification to obtain an overall field of view. The staining was considered of good quality when well-labeled, dark silver impregnated fibers were homogenously distributed across the sections. Scores indicated: Absence=0 (Figure 2K), Heterogenous=1 (Figure 2L), Homogenous=2 (Figure 2M).

##### Background

Silver stains are known to use toxic chemicals and are capricious, unpredictable and difficult to perform (Litchfield & Nagy, 2001; Ogawa, 1990), since silver binds and stains to chemicals, slides, mounting medium, dust, etc. Therefore, the background level of staining was assessed since non-specific silver deposits could vary according to the fixative solution and other variable conditions. Therefore, we observed the presence or absence of silver deposits in the background using the 60X objective, in which the presence of silver spots decreases the contrast of the well-stained fibers against the background, and therefore, a poorer quality of this staining. Scores indicated: Dark=0 (Figure 2N), Intermediate=1 (Figure 2O), Light=2 (Figure 2P).

### 2.7 Confounding variables

All variables mentioned above were also assessed in relation to two confounding variables: the post-mortem interval and the histology delay, to determine whether they might have affected the variables of interest. Raw data is shown in table 1.

#### 2.7.1 Post-mortem interval

The post-mortem interval (PMI) is the time elapsed between death of the donor and the administration of the fixative to the brain. The post-mortem interval could have an impact on the tissue quality since the brain could start to decompose prior to fixation. The acceptable post-mortem interval is 0 to 48 hours, although in cases where the bodies have been refrigerated the interval may be extended an additional 24 hours. To assess this variable, we created two categories, for shorter or longer PMI: Short = 0-24 hours; Long = >24 hours.

#### 2.7.2 Histology processing delay

The histology delay (HD) corresponds to the time elapsed between the moment the brain was fixed (by perfusion in the anatomy laboratory), or from the moment it was immersed in NBF (brain bank technique), to the moment the histology procedures began. Since we used a convenience sample, this variable was not controlled for all specimens. Some brain blocks were obtained after the brains were used for years for teaching (Anatomy laboratory, UQTR (and therefore immersed in 10% ethanol); immediately after the brains were extracted (if fixed by perfusion, UQTR), or after years of immersion in NBF (Douglas-Bell Canada brain bank). These differences may affect the brains differently by causing different levels of cross-linking of the proteins (i.e., antigens). Therefore, we assessed this confounding variable creating two categories, for shorter or longer HD: Short = 0-2 years; Long = >2 years.

### 2.8 Statistical analyses

All categorical variables described above were compared across the three fixative groups using Chi-square tests. All statistical analyses were adjusted for multiple comparisons using Bonferroni correction and were performed using SPSS statistics software (28.0.0 version).

To assess for confounding variables, we also used Chi-square tests to relate the variables of interest (section 2.6) to each confounding variable (section 2.7), first considering all specimens together, and then separately, within each fixative group (NBF; SSS and AFS).

## 3. Results

### 3.1 Ease of manipulation

We found no statistically significant differences between treatment groups in the ease of manipulation scores, but we observed more cases fixed with SSS that scored poor for this criterion, while brains fixed with NBF never scored very poor for ease of manipulation. Also, a higher proportion of cases fixed with NBF and AFS scored good for this criterion (Figure 3).

**Figure 3:**
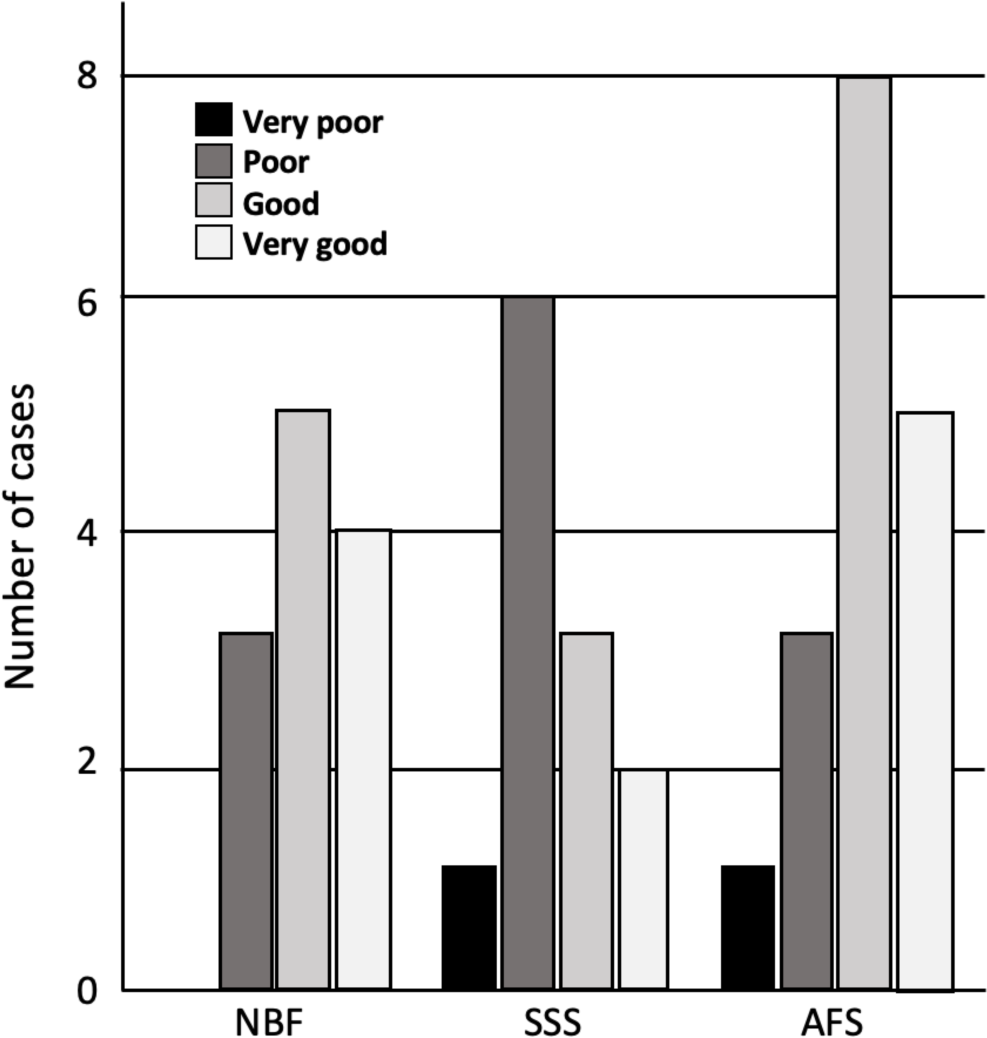
Bar charts of the ease of manipulation of the tissue. Very poor (black)=4 signs; Poor (dark gray)=3 signs; Good (light gray)=1 or 2 signs; Very good (white)=no sign; NBF=neutral-buffered formalin; SSS=salt-saturated solution; AFS=alcohol-formaldehyde solution.

### 3.2 Tissue quality

Tissue quality was assessed according to two criteria, namely the uniformity of the neuropil and cellular shape. We found no significant difference in the frequency of occurrence of fissured neuropil across the three groups (Figure 4A). However, we found a significant difference in the frequency of scores for the cellular shape, between the three groups (p<0.001). Regular cell contours were always observed in the NBF-fixed brains (p<0.001), while irregular (shrunken and shriveled) cell contours were always found in SSS-fixed brains (p<0.001) (Figure 4B).

**Figure 4:**
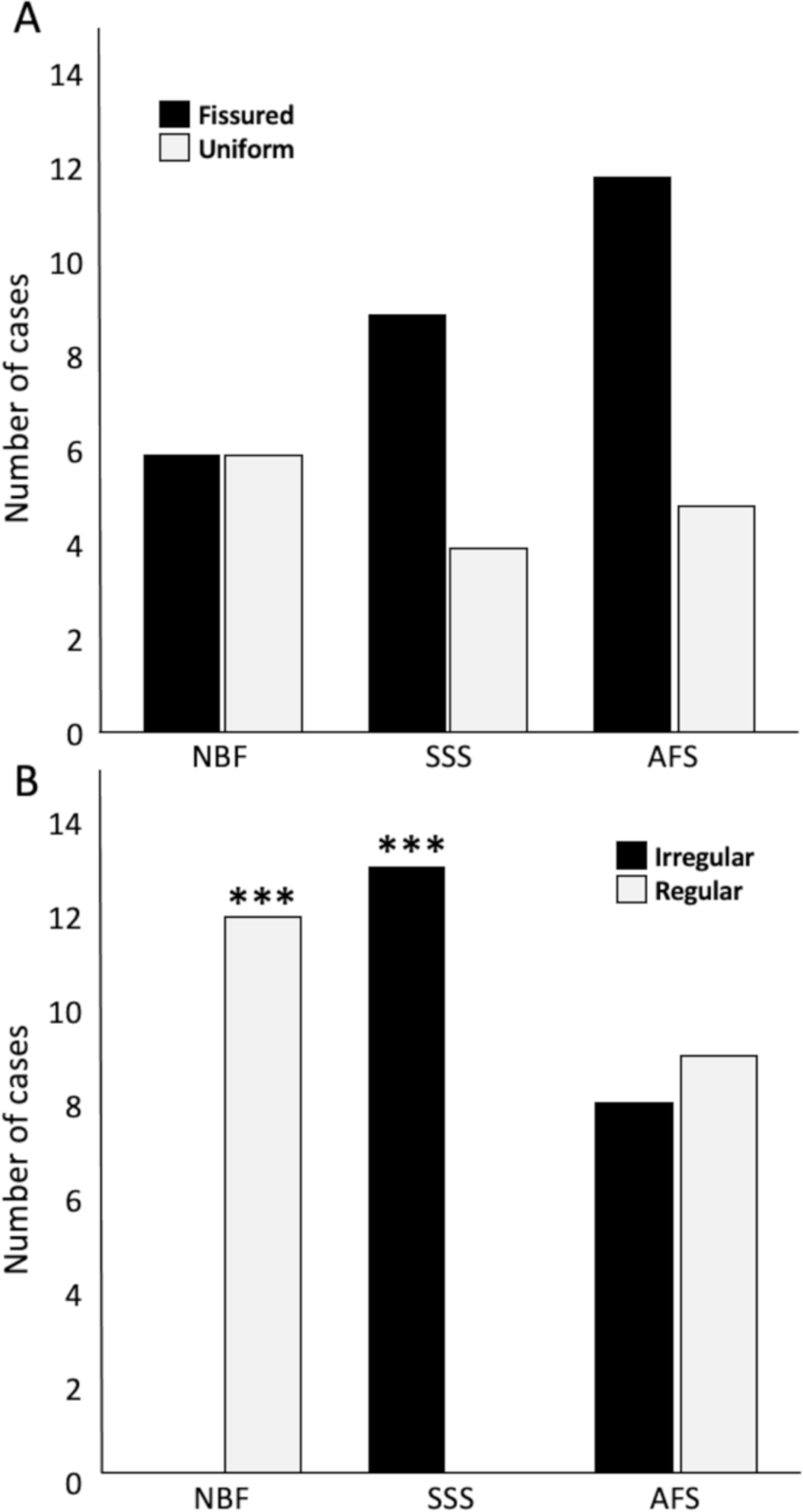
Bar charts of the tissue quality. Neuropil uniformity (A), either fissured=0 (black) or uniform=1 (white), and cellular shape (B), either irregular (shrunken and shriveled)=0 (black) or regular (smooth)=1 (white). ***=Significant after Bonferroni correction (p<0.001); NBF=neutral-buffered formalin; SSS=salt-saturated solution; AFS=alcohol-formaldehyde solution.

### 3.3 Antigenicity preservation

The distribution of labeled cells of the four tested antigens was used to assess antigenicity preservation. A homogenous distribution of NeuN labeling was significantly more frequent in AFS-fixed brains (p=0.0003), whereas significantly higher occurrence of isolated labeled cells (p=0.0019) and no homogenous NeuN labeling (p=0.0019) were observed in SSS fixed brains. NeuN antigenicity was preserved in most NBF-fixed specimens, except for three cases in which no labeled neurons were observed (p=0.005; significant before a Bonferroni correction) (Figure 5A). Homogeneous distribution of GFAP labeling was observed in only one NBF-fixed specimen, and no labeled astrocytes were observed in three cases (p=0.005; significant before a Bonferroni correction) (Figure 5B). Patchy Iba1 labeling occurred significantly more frequently (p=0.0037) whereas homogeneous labeling of this marker occurred less frequently in NBF-fixed brains (p=0.005; significant before a Bonferroni correction). Homogeneous distribution of Iba1 labeling was significantly more frequent, whereas patchy labeling was significantly less frequent in AFS-fixed brains (p<0.001) (Figure 5C). Finally, PLP labeling was more frequently uniform (homogeneous or cell patches) in all AFS-fixed brains (more cases of cell patches: p=0.046 before a Bonferroni correction), Conversely PLP labeling was absent in three SSS-fixed cases and two NBF-fixed cases (Figure 5D).

**Figure 5:**
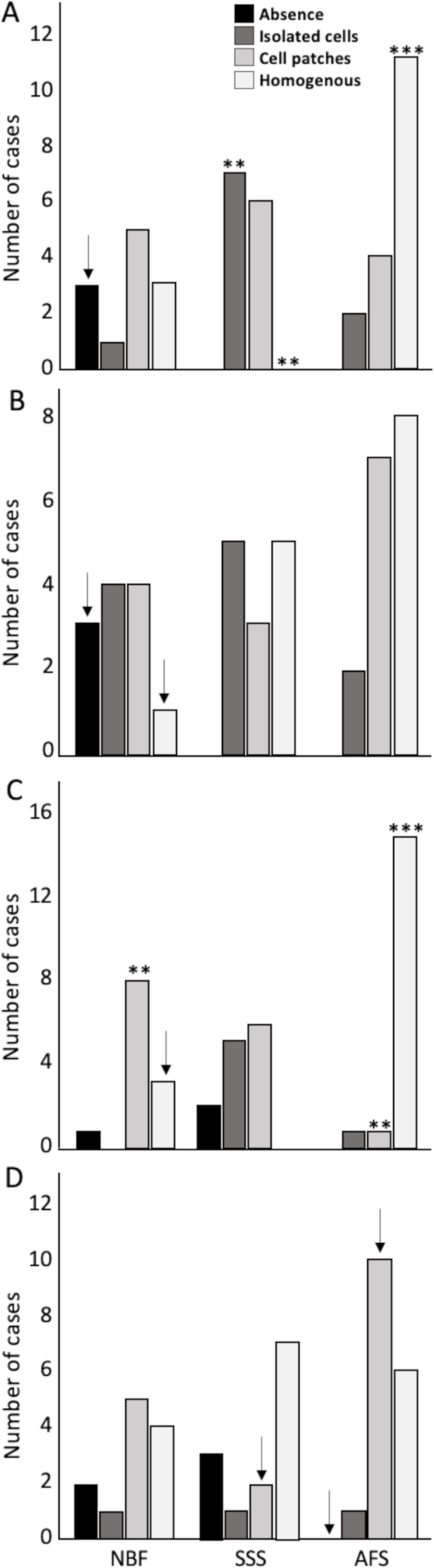
Antigenicity preservation and cellular distribution of the four main brain cell populations. Cellular distribution of Neuronal nuclei (NeuN) (A), astrocytes (GFAP) (B), microglia (Iba1) (C) and oligodendrocytic myelin fibers (PLP) (D). No antigenicity=0 (black); Isolated cells=1 (dark gray); Cell patches= 2 (light gray); Homogeneous distribution=3 (white). **=Significant after Bonferroni correction (p<0.01); ***=Significant after Bonferroni correction (p<0.001); ↓=Significant before Bonferroni correction; NBF=neutral-buffered formalin; SSS=salt-saturated solution; AFS=alcohol-formaldehyde solution.

In cases where our antibodies produced no labeling or only isolated labeled cells, we repeated the IHC procedures following the application of an antigen retrieval protocol. Antigenicity was recovered and produced a homogeneous labeling for all four markers in SSS and AFS-fixed brains, (Figure 6). Nonetheless, NeuN and GFAP antigenicity was not recovered in three NBF fixed cases.

**Figure 6:**
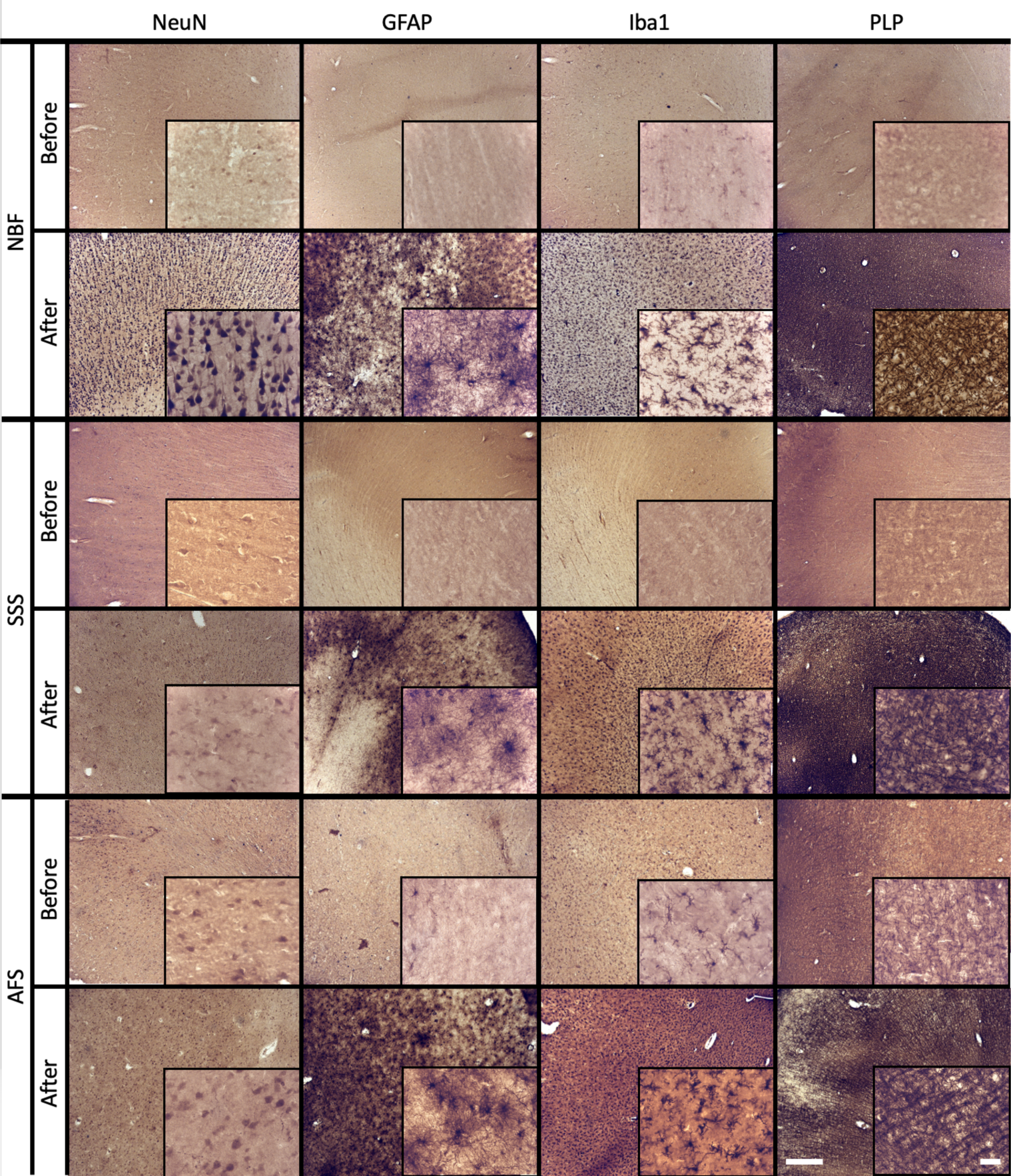
Before and after an antigen retrieval protocol in sections that showed poor antigenicity preservation. Photomicrographs (4X) (inserts are zoomed-in at 40X to see the cells) of cellular distribution of NBF-fixed cases (top), SSS-fixed cases (middle) and AFS-fixed cases (bottom line) that showed no antigenicity (before lines), while a homogenous distribution is shown after an antigen retrieval protocol (after lines). Left scale bar is valid in all 4X photomicrographs = 500 μm. Right scale bar is valid in all inserts (40X) = 50 μm. NBF=neutral-buffered formalin; SSS=salt-saturated solution; AFS=alcohol-formaldehyde solution. NeuN=neuronal nuclei (neurons); GFAP=glial fibrillary acidic protein (astrocytes); Iba1=ionized binding adaptor molecule 1 (microglia); PLP=proteolipid protein (oligodendrocytic myelin).

### 3.4 Histochemical stains

First, we assessed quality of the Nissl-stain. We found a significantly higher frequency of cases showing high cellular contrast in NBF-fixed specimens (strong labeling of neurons in all the cases) (p=0.0037), and less cases with pale neurons (p=0.016; significant before a Bonferroni correction only). The brains fixed with the other two solutions sometimes produced pale neurons staining (SSS: N=6, p=0.045; significant before a Bonferroni correction only, and AFS: N=5), or even no staining at all (SSS: N=1; AFS: N=2) (Figure 8A).

**Figure 7:**
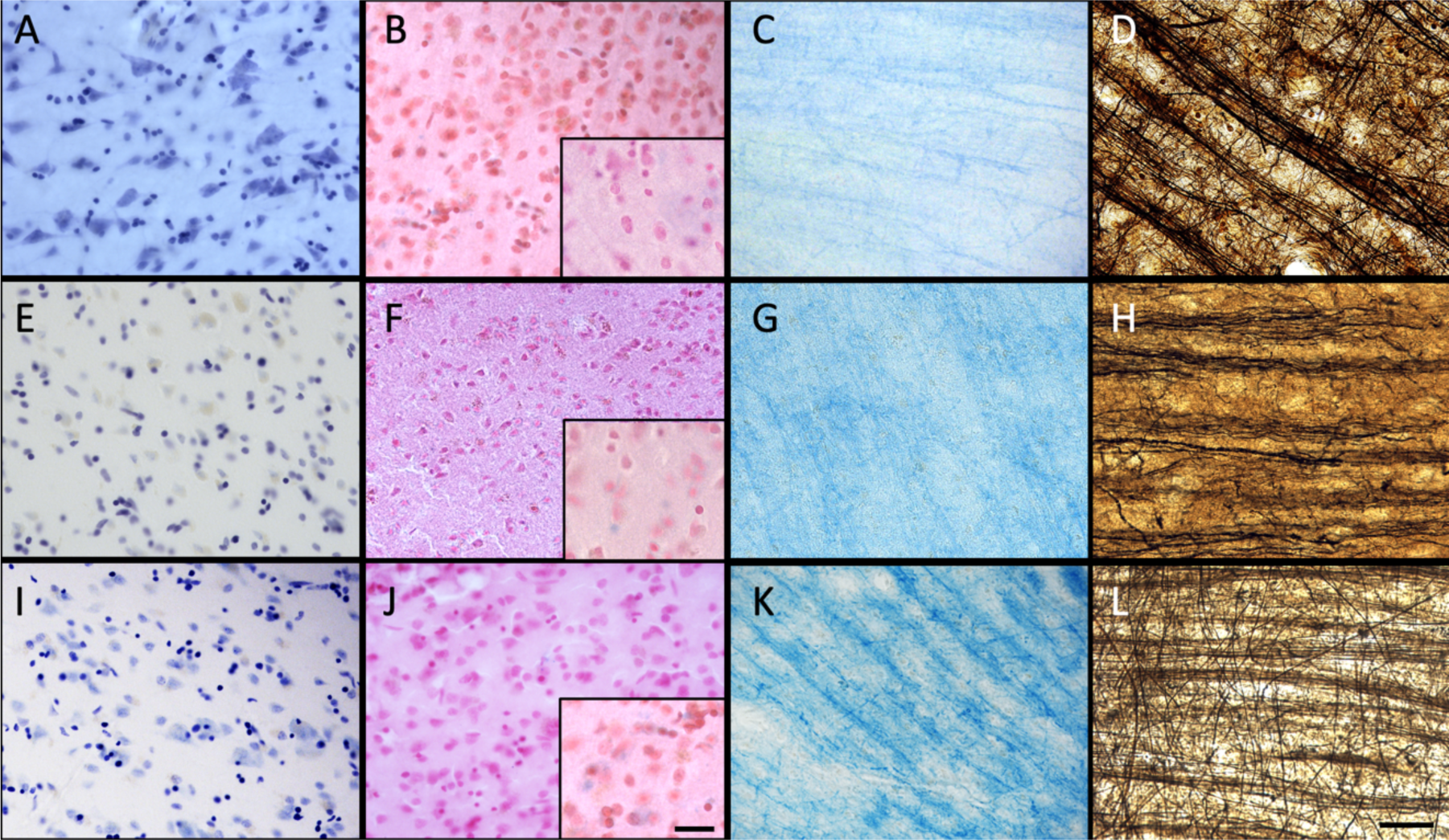
Quality of the histochemical stains. Photomicrographs (40X) of the Cresyl violet staining (neurons and glial cells in dark purple) for NBF (A), SSS (E), and AFS (I), of the Prussian blue staining (neurons and glial cells in pink, iron in blue, with iron showed in inserts (100X) Scale bar = 50μm) for NBF (B), SSS (F), and AFS (J), of the LFB staining (myelin fibers in dark blue) for NBF (C), SSS (G), and AFS (K), and Bielchowsky’s staining (axons and neurofibrils in dark brown) for NBF (D), SSS (H), and AFS (L). Scale bar = 50 μm (valid for A to L).

**Figure 8:**
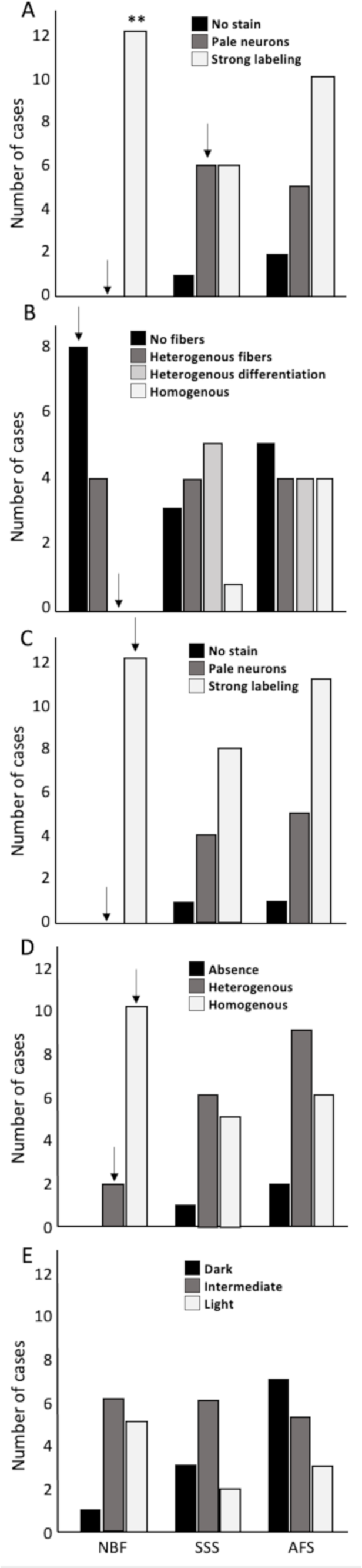
Bar charts of the quality of the histochemical stains. A) Cresyl violet staining quality assessed by the absence of labeling=0 (black), pale neurons=1 (dark gray) or strong labeling=2 (white). B) Luxol fast blue staining quality, assessed by the absence of fibers=0 (black), heterogenous fibers=1 (dark gray), heterogeneous differentiation=2 (light gray) and homogenous staining=3 (white). C) Prussian blue staining quality assessed by the absence of labeling=0 (black), pale neurons=1 (dark gray) or strong labeling=2 (white). D) Fibers preservation in Bielchowsky staining assessed by absence of fibers=0 (black), heterogeneous distribution of the fibers=1 (dark gray), or homogenous distribution=2 (white). E) Bielchowsky’s staining background which could be dark=0 (black), intermediate=1 (dark gray) or light=2 (white). **=Significant after Bonferroni correction (p<0.01); τ=Significant before Bonferroni correction; NBF=neutral-buffered formalin; SSS=salt-saturated solution; AFS=alcohol-formaldehyde solution.

Second, we assessed the quality of myelinated fibers labeling with LFB. NBF-fixed brains showed a higher frequency of cases in which stained fibers were absent (score=0) (p=0.016; significant before a Bonferroni correction only) and never homogeneous distribution (p=0.036; significant before a Bonferroni correction only). Fiber labeling with LFB in SSS and AFS-fixed brains was quite inconsistent as they showed randomly distributed scores (Figure 8B).

Cell to background contrast was high following a Nuclear fast red (Nissl) counterstain to the Prussian blue staining (Figure 8C), regardless of the fixative. High cellular contrast occurred at a higher frequency in NBF specimens (all cases showed dark neurons: score=2) (p=0.016; significant before a Bonferroni correction only). Also, despite that not all the specimens showed ferric iron aggregation (depending on the pathology or age of the patient), blue spots indicating iron deposits were present in 10 NBF, 6 SSS and 8 AFS-fixed brains demonstrating the likelihood of detecting these deposits when either of these fixation protocols are applied (Figure 7, inserts in B, F, J).

Finally, we found a higher frequency of cases showing homogeneous labeling of silver impregnation of axonal fibers with the Bielchowsky procedure in the NBF-fixed brains (p=0.0093; significant before a Bonferroni correction only). However, a higher occurrence of dark or intermediate background staining, was observed in the SSS and AFS compared to the NBF fixed brains, but this difference was not statistically significant.

### 3.5 Confounding variables

#### 3.5.1 Post-mortem interval

More than half of the specimens were harvested following a short PMI (25 of 42). Longer PMI occurred more often in the NBF-fixed group than the other two groups (8 of 12; p=0.028; significant before a Bonferroni correction only).

Ease of manipulation scores were very good in more cases with a shorter PMI (p=0.016; significant before a Bonferroni correction only), whereas longer PMI cases had a higher frequency of good scores (p=0.012; significant before a Bonferroni correction only). More specimens with shorter PMI were very easy to manipulate in brains fixed with NBF, (p=0.028), while specimens with longer PMI only scored good or poor for this criterion. This could indicate better tissue preservation following short PMIs; however, statistical significance difference did not resist the Bonferroni correction.

PMI did not significantly affect quality of tissue and of neuropil scores. However, we did observe more broken and less uniform neuropil in cases with both shorter (15 out of 25) and longer (12 out of 17) PMI. This slight, albeit non-significant, advantage of short PMI however does not translate to tissue quality scores. Indeed, 15 specimens with a shorter PMI showed irregular cells, compared to 10 cases in which regular cells shapes were observed, which would suggest an unexpected disadvantage for shorter PMI, but we had 6 specimens with longer PMI that showed regular contours, and 11 showed irregular cell contours, suggesting otherwise.

The effect of the PMI on the preservation of antigenicity was assessed separately for the four antibodies. For NeuN, we found that the cases that showed no cell labeling were in the longer PMI group, which was statistically significant before a Bonferroni correction (p=0.028). Furthermore, we found more cases with a homogenous or patchy cell distribution in cases with a shorter PMI, but this was not statistically significant. When assessed within each fixative group, we found unlabeled cells more often in NBF-fixed brains with longer PMI, and homogeneous patchy distribution of cells in the cases with a shorter PMI, although this observation did not reach statistical significance. We found that there were more cases with homogenous distribution and fewer specimens with no GFAP labeling when fixed within a shorter PMI, but there were no statistical labeling patterns in cases with a shorter or longer PMI. This contrasts with the AFS-fixed brains where homogenous GFAP labeling was found in eight specimens with short PMI, compared to none with a longer PMI (statistically significant prior to Bonferroni correction: p=0.012). We found no statistical difference of Iba1 labeling between the two groups (short or long PMI), and this was still true when assessing within each group. Similarly, no statistical difference was observed for the two groups for PLP labeling. However, in NBF-fixed brains, we found that there were more cases with a short PMI (N=2) with no PLP labeling, while no such cases were observed in the group with a longer PMI, although this did not maintain significance after a Bonferroni correction (p=0.028).

We also assessed the quality of histochemical stains of the specimens in relation to the shorter or longer PMI. We found no statistical differences of the quality of Cresyl Violet, Luxol fast blue and Prussian blue stains between the groups. However, fiber labeling with the Bielchowsky stain was more often heterogeneous in cases with a shorter than with a longer PMI (p=0.036), although this did not retain significance after Bonferroni correction. There were no significant differences in the fiber labeling within group. The quality of the background labeling of the Bielchowsky’s stain was not different between the PMI groups or fixatives.

#### 3.5.2 Histology delay

The occurrence of short and long HDs was not statistically significant between the fixative groups; 20 specimens showed shorter and 22 showed longer delays.

We found no statistical difference in the ease of manipulation of sections between cases with a shorter or longer histology delay (HD). This observation was also true when comparing the shorter or longer delays within the fixative group.

We found no significant difference of the neuropil uniformity between the cases with a shorter or longer HD, with most specimens showing a broken neuropil in both groups. This was also observed when assessing within group. There was no difference of the cellular shape scores between cases with shorter or longer HD for SSS and NBF-fixed cases. For AFS, although not statistically significant, we observed a higher occurrence of irregular (shrunken and shriveled) cells in cases with longer HD (N=7), while only one case with irregular cells was observed with shorter HD.

We found no significant difference distribution of NeuN labeled cells between the shorter or longer HD although the only specimens that showed no labeled cells had a longer HD (N=3) and were fixed with NBF. We found no differences when comparing the HD within each fixative group. We found significantly more cases with a homogenous distribution of GFAP labeling in cases with a shorter HD (p=0.0051), which was related to the AFS-fixed brains that showed more homogenous distribution in the shorter HD (p=0.028), and a patchier distribution of labeled cells with a longer HD (p=0.012), although this did not remain significant after a Bonferroni correction. Furthermore, although not statistically significant, homogenous distribution of GFAP labeling was never found in cases with longer HD following both SSS and NBF fixation. For Iba1 and PLP, we found no statistical difference between the two groups.

We also assessed the quality of histochemical procedures in relation to HD length. We found no statistical differences in the Cresyl Violet, Luxol fast blue and Prussian blue staining quality between the groups. However, we found a higher frequency of unlabeled and faintly labeled cells by Cresyl violet and the Nuclear red counterstaining, in AFS-fixed brains with a long HD, but this did differences did not reach statistical significance. We found no statistical difference between shorter or longer HD overall for the quality of Bielchowsky silver impregnation of nerve fibers. However, when assessing within fixative group, we found that the labeled fibers were more homogeneously distributed in NBF-fixed brains, in both groups (shorter and longer HD), while SSS-fixed brains with longer HD showed absence or heterogenous fiber labeling, but this did not reach statistical significance. Finally, in the AFS-fixed brains, no labeled fibers were observed in the shorter HD group (p=0.046; significant before a Bonferroni correction only). The Bielchowsky’s background was not affected by the histology delay. However, a higher frequency of cases with light backgrounds was found in the NBF-fixed brains with shorter HD and more cases with intermediate backgrounds with longer HD (p=0.021, significant before a Bonferroni correction only). We also found more cases with dark Bielchowsky backgrounds with longer HD in AFS-fixed brains, but this was not statistically significant.

## 4. Discussion

### 4.1 Ease of manipulation

We were expecting that the SSS and AFS-fixed brains would be difficult to manipulate (softer tissue), according to our previous study in mice (Frigon et al., 2022). Accordingly, recent study in rat brains (Martins-Costa et al., 2022), and human anatomical studies describe these fixatives as soft-embalming solutions, which could led to differences in the texture of the brains (Balta et al., 2015; Eltoum et al., 2001; Hammer et al., 2012; Hayashi et al., 2014; Hopwood et al., 1989; Nicholson et al., 2005; Rahman et al., 2021; Tomalty et al., 2019). The human tissue samples were relatively stiff and well-fixed macroscopically, and we found mostly that sections from AFS fixed specimens were easy to manipulate, and slightly more difficult in cases fixed with SSS. Nevertheless, SSS-fixed human sections were relatively easier to manipulate (stiffer) than the mouse sections of brains fixed with the same solutions (Frigon & al., 2022). This could be related to their different capacity to absorb the fixative (diffusion), related to their differences in shape, size, gyrification, and vascularization circuits across species (Ventura-Antunes et al., 2013).

### 4.2 Tissue quality

Integrity of the neuropil was compromised in the three groups in which we found a higher occurrence of fissures. Conversely, a more uniform neuropil was observed in NBF-fixed mouse brain sections (Frigon et al., 2022). This could be explained by the quick perfusion fixation of mice and timely histological processing soon after fixation compared to the lengthy months to years delay before histological processing of the brain bank NBF-fixed human specimens which may enhance chemical artefacts on the tissue. Regarding the broken neuropils, the high concentration of isopropyl alcohol or ethanol in SSS- and AFS-fixed brains, likely dehydrates tissues significantly (Viktorov & Proshin, 2003), and may cause fissures in the neuropil by shrinking, by changing the osmolarity of the intracellular vs. extracellular spaces. This may also explain the irregular aspect of the cells in the SSS and AFS-fixed brains, compared to the NBF-fixed brains that only show regular cell contours.

### 4.3 Antigenicity preservation

We found that NeuN and GFAP antigenicity was not well preserved in all cases, especially in NBF-fixed brains, where no cells were observed in three specimens. This was expected of the NBF-fixed brain bank specimens, which are preserved in a solution with a higher concentration of formaldehyde often for years before histology processing (i.e., longer histology delays). The cross-linking, due to the over-fixation by formaldehyde immersion (Arnold et al., 1996; Stumptner et al., 2019) likely affected the conformation of the neuronal nuclei and the glial fibrillary acidic protein more profoundly than the other two analyzed antigens, due to a higher interaction between their specific tertiary conformations and the chemicals, which is in line with the qualitative description of formaldehyde and its chemical reactions (Fox et al., 1985; Helander, 1994; Kiernan, 2000; Puchtler & Meloan, 1985; Thavarajah et al., 2012). Moreover, an antigen retrieval protocol recovered the antigenicity for Iba1 and PLP in all cases in a homogeneous way, whereas NeuN and GFAP did not regain antigenicity in some NBF-fixed brains.

Antigenicity was generally best preserved by the AFS fixative. SSS fixation produced mostly labeling of isolated cells or patches. However, an antigen retrieval protocol recovered all the antigens in tissue fixed with the two solutions used in anatomy laboratories, either SSS or AFS. Therefore, sufficient quality IHC procedures are possible following the application of an antigen retrieval protocol. These observations differ from what we reported in our previous study in mice, in which the antigenicity of the four main cell populations was preserved and homogeneously distributed across all the types of fixation (Frigon & al., 2022), even when no antigen retrieval protocol was applied prior to the IHC procedures. This also supports the differences in mice and human brain tissue preservation properties.

### 4.4 Histochemical stains

The clear labeling of cells with Cresyl violet and the Nuclear red counterstain of the Prussian blue procedure, demonstrates that the brains fixed with solutions from anatomy laboratories could be used for cytoarchitectonic analysis (Amunts et al., 2020; Amunts & Zilles, 2015; García-Cabezas et al., 2020), brain morphology (Zachlod et al., 2022), neurodegenerative disease (Bobinski et al., 2000), and MRI validation (Alkemade et al., 2020; Hoffmann et al., 2011; Magnain et al., 2014). Furthermore, Prussian blue (Perl’s stain) was feasible in specimens fixed with the three solution and helped detecting ferric iron deposits.

Luxol fast blue fiber staining was not homogenous in NBF fixed brain sections. In some NBF fixed cases, labeling luxol fast blue failed to stain myelinated fibers altogether. Similarly, luxol fast blue staining of myelinated fibers was not consistent in both SSS and AFS-fixed brains suggesting that IHC (i.e., PLP in our case) might be a better approach to routinely label the myelinated axons. Indeed, IHC (i.e. MBP, MAG, PLP antigens) are widely used in neuropathology (Adiele & Adiele, 2019; Beach et al., 2023; Kuhlmann et al., 2017; Stüber et al., 2014; Vakrakou et al., 2022).

Bielchowsky silver impregnation of fibers, was best achieved in NBF-fixed brains, followed by SSS and AFS in which more cases with heterogeneous staining and intermediate density backgrounds were observed. This may be related to the ethanol and/or amount of alcohols in their chemical compositions. An old study on silver impregnation technique showed that lipid extraction with organic solvents such as ethanol can lead to unsatisfactory impregnation and poor staining (Wolman, 1958). However, we found good cases and high-quality stains following all fixation procedures, suggesting that the fixatives used in anatomy labs may be reliable for neuroscientific research.

### 4.5 Confounding variables

It is known that molecular conformations change after death, at different paces depending on the molecules being analyzed, for some cases within minutes or the first few hours of post-mortem interval before fixation. We therefore expected better histological results in brains with shorter PMI (Beach et al., 2015; Hynd et al., 2003; Palmer et al., 1988; Ravid et al., 1992; Spokes, 1979) and with a short HD, avoiding post-mortem tissue decay and excessive cross-linking from over-fixation (Arnold et al., 1996; Schulz et al., 2011; Stumptner et al., 2019). Surprisingly, we found no evidence that neither lengthy PMI nor HD affected the tissue more significantly than the fixative itself; some quality histology was obtained even in the cases with longer delays, and some poorer cases were observed in the shorter delays. It is worth noting that some individual cases suggested a slight influence of the PMI in the NBF-fixed brains, on the ease of manipulation, and antigenicity preservation of NeuN. These NBF brain bank tissue samples seemed the most impacted by the PMI, since they had significantly greater PMI (median = 45 hours) than the anatomy laboratory specimens (median = 22 hours) (p<0.001). Finally, we found significantly more homogenous GFAP distribution in cases with a shorter HD and PMI in AFS-fixed brains, suggesting that this antigen might be more affected by these confounding variables. Other results were not impacted in AFS-fixed brains, in which all histological procedures were successful.

Finally, to ensure that our interpretation of the confounding variables was sufficiently robust, we also ran a multinomial logistic regression for each variable of interest, finding overall similar results. Only the observation of AFS-fixed brains with a long HD that showed a higher frequency of pale or absent cells using Nissl Stains that was previously non-significant did became significant (p=0.008) when calculating the results using this regression model.

Our results suggest that overall PMI and HD do not seem to impact the tissue quality. However, we did have a bit of an impact in some specific variables, such as NeuN antigenicity that was more frequently absent in long PMI in NBF-fixed brains (albeit no statistically significant). So, we consider that the prioritization of shorter delays should be preferred when feasible. Future studies increasing the number of specimens will elucidate more precisely the impact of longer delays on specific antigens.

### 4.6 Anatomy laboratory’s fixatives

#### 4.6.1 Salt saturated solution

We suggest that the SSS-fixed brains could be appropriate for neuroscientific research, since all the procedures were successful. However, these tissue blocks were more difficult to manipulate, and the overall quality of the tissue was poorer, thus SSS-fixed brains require more careful manipulation. Furthermore, antigenicity preservation was inadequate, but we demonstrated that IHC is still feasible following the application of an antigen retrieval protocol. We speculate that the lighter fixation of SSS brains owing to the low formaldehyde concentration led to a poor tissue preservation. The high osmolarity of this solution might also impact the diffusion of the fixative through the capillaries and interstitial compartments of the brain. Indeed, the SSS is over saturated in salt, which could lead to absorption of the free water in the brain prior to fixation with phenol and the small amount of formaldehyde contained in the solution (Coleman & Kogan, 1998; Hayashi et al., 2014), generating a lighter fixation. In addition, the cryoprotection in sucrose was very long (i.e., over a week compared to 1-2 days for the blocks fixed with the other two solutions) for these specimens; this might have also affected the tissue quality creating freezing artifacts.

#### 4.6.2 Alcohol-formaldehyde solution

Our results show that the AFS-fixed brains provided the best quality histology. Tissue was manipulated with ease, was of good quality, and produced the best IHC antigenicity preservation and quality of histochemical stains. Here again, an antigen retrieval protocol prior to the IHC procedures was warranted for optimal results in some specimens. The brains from the AFS group were also, for the most part, the oldest specimens (with a range of HD between 1 and 15 years), but even the old specimens held up well. AFS contains a higher concentration of formaldehyde (1.4%), compared to SSS (0.3% formaldehyde), which probably improved the fixation level of the brain. On the other hand, the lower formaldehyde concentration in the AFS compared to NBF, might reduce excessive cross-linking, making for the better preservation of antigenicity. Therefore, our results suggest that fixation with approximately 1% formaldehyde might be ideal and should be tested in future studies.

It is known that alcohol solutions might be optimal for combining gross anatomy dissection, surgical training and histology assessment of different tissues (Barton et al., 2009; Benet et al., 2014; Hopwood et al., 1989; Martins-Costa et al., 2022; O’Sullivan & Mitchell, 1993; Rahman et al., 2021; Tomalty et al., 2019; Venne et al., 2020). In this work, we have extensively assessed our specific solution, as it differed, although slightly, from the other formulas that are used worldwide (Barton et al., 2009; Benet et al., 2014; Cabrera et al., 2017; Hopwood et al., 1989; Martins-Costa et al., 2022; O’Sullivan & Mitchell, 1993; Rahman et al., 2021; Shetty et al., 2020; Tomalty et al., 2019; Venne et al., 2020). To our knowledge, no brain histology quality and antigenicity preservation assessments are available in the literature for such solutions. Altogether, our results suggest that AFS-fixed brains are of sufficient quality for qualitative or/and quantitative neuroscientific research.

### 4.7 Study limitations

First, working with human post-mortem tissue involves dealing with confounding variables, such as the post-mortem delay and the histology delay. The brains could also be affected by other variables such as the age, sex or presence of comorbidities. The post-mortem and histology delays have been assessed as confounding variables, and we found that they did not significantly impact our results. Although our specimens were from a convenience sample, they mostly included normal-aging brains with absence of neuronal or vascular comorbidities, with a random distribution of sex. Second, using a convenience sample ensured that we obtained tissues from different brain lobes. However, we confirm that we used neocortex for all the specimens. Third, free-floating tissue sections stained with IHC procedures might show heterogeneous labeling, since the staining depends on the level of antibody penetration and the tissue background. Analyzes might therefore be affected by the protocol itself, which was also evaluated as a variable of interest in our previous study in mice brains (Frigon & al., 2022). Therefore, the protocols were already tested and validated for this study, so we excluded the antibody penetration and tissue background quality as variables here. Fourth, we are aware that other antigens of interest could have been tested, but we limited our study to the antigens that were previously validated in our study in mice (Frigon & al., 2022). Finally, the statistical power might have been limited by our relatively small sample size, especially when dealing with confounding variables. However, availability of human brain tissue is limited, and our study included all the specimens that were available at the time of the assessments. Future studies with larger sample sizes are warranted to further our understanding of the impact of confounding factors.

### 4.8 Conclusion

In this study, we compared the quality of histochemical and immunohistochemical procedures performed on brain tissue from cases fixed by current embalming protocols in use in gross anatomy dissection laboratories and fixation protocols in use in major brain bank settings. We showed lower overall histology quality using a salt saturated solution but also a higher quality for some specific variables when using an alcohol-formaldehyde solution in comparison to the classic neutral-buffered formalin solution used in brain banks. We further showed that immunohistochemistry and histochemical stains are feasible with brains fixed with two solutions used in gross anatomy laboratories following the application of a heat-induced antigen retrieval protocol. Therefore, brains coming from body donation programs for teaching gross-anatomy could also be used for neuroscientific research, which could increase the amount of brain tissue available to neuroscientists.

## Conflict of interest

The authors declare that the research was conducted in the absence of any commercial or financial relationships that could be construed as a potential conflict of interest.

## Author contribution

DB and JM contributed to the conception and design of the study. EMF performed the experimental protocols, organized the database and performed the statistical analysis. AGL performed a part of the experimental protocols. EMF wrote the article. MD contributed with samples for data generation. JM and DB wrote sections of the article, and they contributed to manuscript revision with MD. All authors read and approved the submitted version.

## Funding

Gouvernement du Canada | Natural Sciences and Engineering Research Council of Canada (NSERC):

Eve-Marie Frigon BESCD3-559015-2021

Denis Boire RGPIN-2018-06506

Josefina Maranzano DGECR-2021-00228

## Acknowledgements

We would like to thank the incredibly generous body donators and their family for making our project possible. We also acknowledge funding resources and the Anatomy Laboratory staff (Johanne Pellerin, Marie-Eve Lemire, Sonia Gauthier and Sophie Plante) of the University of Quebec in Trois-Rivieres for their support and help at the lab.

## Notes

### Competing Interest Statement

The authors have declared no competing interest.

